# A universal plug-and-display vaccine platform for mucosal and systemic immunity using *Bacillus* subtilis membrane vesicles

**DOI:** 10.64898/2026.08.12.744535

**Authors:** Kimihiro Abe, Takehito Wakabayashi, Hiroki Kawabata, Kozue Sato, Ryoma Nakao, Takehiro Yamaguchi, Hirotaka Kobayashi, Michiyo Kataoka, Tsutomu Sato, Yukihiro Akeda

**Affiliations:** Department of Bacteriology I, National Institute of Infectious Diseases, Japan Institute for Health Security (JIHS), Shinjuku-ku, Tokyo, Japan; Research Center for Vaccine Development, National Institute of Infectious Diseases, Japan Institute for Health Security (JIHS), Shinjuku-ku, Tokyo, Japan; Department of Frontier Bioscience, Hosei University, Koganei-shi, Tokyo, Japan; Department of Infectious Disease Pathology, National Institute of Infectious Diseases, Japan Institute for Health Security (JIHS), Shinjuku-ku, Tokyo, Japan

**Keywords:** *Bacillus subtilis*, bacterial membrane vesicle, vaccine, plague, *Yersinia pestis*

## Abstract

Many bacterial species naturally secrete membrane vesicles (MVs) that mediate the intercellular transport of biomolecules, including nucleic acids, proteins, and metabolites. Beyond their native physiological roles, MVs hold considerable potential for biomedical applications. Here, we demonstrate that MVs from several *Bacillus* species exhibit potent intrinsic adjuvant activity, efficaciously eliciting immune responses and facilitating antigen-specific antibody production in mice. Exploiting this adjuvanticity, we engineered a highly adaptable universal vaccine platform that uses *B. subtilis* MVs as self-adjuvanting carriers. This system employs a modular “plug-and-display” architecture that covalently anchors recombinant antigens to the MV surface through a multi-step bioconjugation cascade. After validation of this methodology using a model antigen, we adapted the platform to target *Yersinia pestis*, the causative agent of plague. We formulated a *Y. pestis* vaccine by labeling the MV surface with a modified capsule antigen fraction 1 (mCaf1). Intranasal administration of the mCaf1-MV vaccine effectively elicited both systemic and mucosal immunity. Crucially, this vaccine conferred highly efficacious protection against a lethal *Y. pestis* infection in a murine model. These findings demonstrate the exceptional protective efficacy of the *B. subtilis* MV platform and highlight its broad potential for the rapid development of mucosal vaccines against diverse emerging pathogens.

## Introduction

Membrane vesicles (MVs) are naturally occurring spherical nanostructures, typically 20−400 nm in diameter, secreted by a wide array of bacterial species. MVs were initially characterized in a Gram-negative bacterium, *Escherichia coli*, which blebs and releases portions of its outer membrane when starved for amino acids (Work *et al*., 1966, Bishop & Work, 1965); hence, those vesicles are termed outer membrane vesicles (OMVs) (Schwechheimer & Kuehn, 2015). Subsequent investigations have established that Gram-positive bacteria also produce MVs derived from the cytoplasmic membrane (termed cytoplasmic membrane vesicles, although referred to herein collectively as MVs). This phenomenon has been documented in many Gram-positive bacteria, including *Staphylococcus aureus* (Wang *et al*., 2018), *Clostridium species* (Obana *et al*., 2017, Kobayashi *et al*., 2022), and *Bacillus subtilis* (Brown *et al*., 2014, Toyofuku *et al*., 2017, Abe *et al*., 2021). Biologically, MVs function as delivery vehicles, encapsulating enzymes, metabolites, signal molecules, and nucleic acids to facilitate biofilm development/maintenance, cell–cell communication, and horizontal gene transfer (Caruana & Walper, 2020, Zhang *et al*., 2026).

In translational medicine, MVs have emerged as an innovative and promising vaccine modality. Because they are non-replicating, immunogenic nanoparticles, they efficiently move into the lymphatic system and are naturally enriched with pathogen-associated molecular patterns (PAMPs), such as lipopolysaccharides (LPS), lipoproteins, peptidoglycans, extracellular nucleic acids, and flagella (Li *et al*., 2026). These PAMPs act as intrinsic self-adjuvants, engaging pattern recognition receptors (PRRs) to heavily stimulate both innate and adaptive immune pathways (Ellis & Kuehn, 2010, Gan *et al*., 2023). Furthermore, intranasal administration of MVs has been shown to effectively elicit dual mucosal and systemic immunity (van der Pol *et al*., 2015, Gerritzen *et al*., 2017). Importantly, mucosal immunity plays a crucial role in blocking pathogens transmitted via air or droplets, as mucosal surfaces are the primary entry points of such pathogens (Zhou *et al*., 2025). Exploiting these attributes, MV-based vaccines have shown promise in inducing both humoral and cell-mediated immunity against various bacterial pathogens, including *Neisseria meningitidis* (Koeberling *et al*., 2009), *Bordetella pertussis* (Bottero et al., 2016), *Pseudomonas aeruginosa* (Zare Banadkoki et al., 2022), *Acinetobacter baumanni* (McConnell *et al*., 2011), and *Vibrio cholerae* (Sedaghat *et al*., 2021). However, the clinical application of pathogen-derived MVs presents formidable challenges. These vesicles often harbor inherent safety risks owing to residual virulence factors and the high endotoxicity of LPSs (Varol *et al*., 2025). Furthermore, the industrial-scale cultivation of bacterial pathogens poses significant biohazard risks, and the strategy of using MVs as vaccines is limited to MV-producing bacterial pathogens.

To circumvent these fundamental limitations, we engineered a universal MV vaccine platform predicated on a modular “plug-and-display” architecture (Figure 1). In this paradigm, MVs from non-pathogenic bacteria serve as universal, self-adjuvanting carriers. Custom vaccines are formulated by labeling the MV surface with target recombinant antigens via a robust covalent linkage cascade, using chemical crosslinkers, click chemistry (Knall & Slugovc, 2013), and DogTag/ DogCatcher-mediated protein ligation (Keeble *et al*., 2022). This strategy profoundly expands the versatility of MV-based vaccines, allowing for the rapid deployment of countermeasures against various pathogens, including non-MV-producing infectious agents such as viruses. As MV vaccine carriers, non-pathogenic, “generally recognized as safe” (GRAS) bacteria are highly biocompatible and therefore ideal sources. In this study, we focused on *Bacillus* species because they lack LPSs in their membrane. Through systematic screening, we established *Bacillus subtilis*, a Gram-positive, non-pathogenic, probiotic bacterium, as our optimal MV-producing host. In contrast to the extensive characterization of MVs from *E. coli* and other Gram-negative bacteria, the immunological potential of Gram-positive bacteria remains largely underexplored, presenting a critical frontier in modern vaccinology.

**Figure 1.**
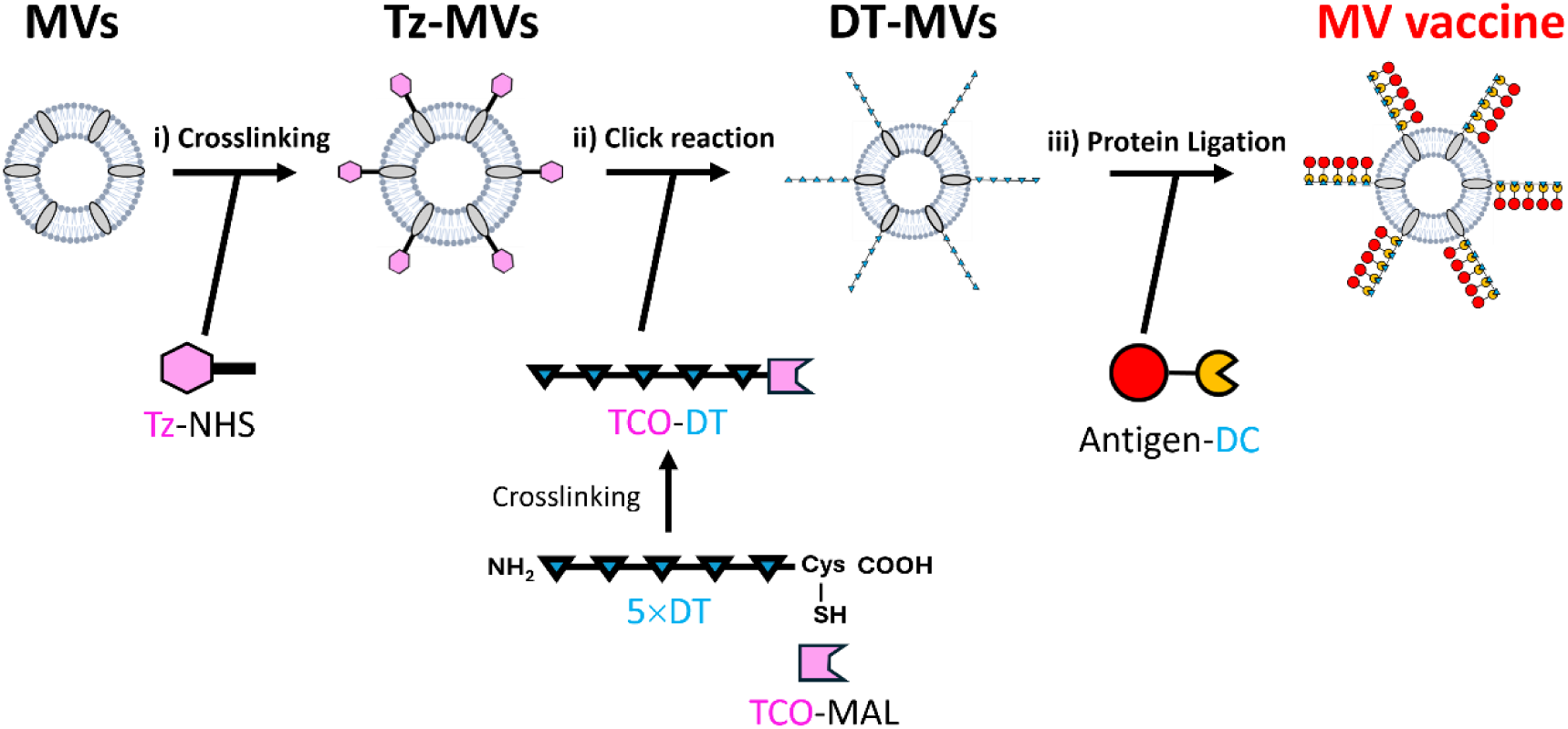
Schematic of MV vaccine production. i) Tetrazine labeling of *B. subtilis* membrane vesicles (MVs). *B. subtilis* MVs isolated from the culture supernatant are reacted with tetrazine N-hydroxysuccinimide ester (Tz-NHS). The MV surface is labeled with Tz through a coupling reaction between NHS and primary amine groups of MV components, yielding Tz-labeled MVs (Tz-MVs). ii) Attachment of the 5×DT scaffold onto Tz-MVs. First, the C-terminal cysteine residue of the scaffold protein containing five DogTag repeats (5×DT) is labeled with *trans*-cyclooctene-PEG3-maleimide (TCO-MAL) to generate TCO-DT. Subsequently, TCO-DT and Tz-MVs are covalently linked via a click reaction between Tz and TCO in PBS, generating 5×DT-attached MVs (DT-MVs). iii) Antigen anchoring onto DT-MVs. Target antigens fused to DogCatcher (DC) are covalently conjugated to the 5×DT on the DT-MVs via spontaneous isopeptide bond formation between DT and DC, yielding the final MV vaccine.

In this study, we first established the bioconjugation methodology of this universal vaccine platform, using a mScarlet-I fluorescent protein as a model antigen. Building on this foundation, we subsequently adapted the platform to target *Yersinia pestis*, developing a highly specialized vaccine by using recombinant *Y. pestis*-specific antigens.

*Y. pestis* is a Gram-negative bacterium and the etiological agent of the plague, remains one of the most lethal pathogens in human history (Perry & Fetherston, 1997). Despite decades of research, no licensed plague vaccine is currently available in the U.S. or Europe (Verma & Tuteja, 2016): production of a heat-killed whole cell (KWC) vaccine was discontinued in 1999 because of severe side effects, and a live attenuated vaccine (LAV) using *Y. pestis* EV76 is fully virulent under iron-overload conditions, *i.e*., in individuals with hemochromatosis (Williamson, 2024). We comprehensively evaluated the immunological response elicited by mCaf1-MVs and definitively demonstrated its protective efficacy against a lethal *Y. pestis* challenge in a murine model. Our findings underscore the profound versatility of the *B. subtilis* MV vaccine platform, offering a highly adaptable and streamlined paradigm for rapid vaccine development.

## Results

### Engineering the universal vaccine platform

As schematically illustrated in Figure 1, we engineered a “plug-and-display” modular vaccine system. By decoupling MV production from antigen expression, this platform enables the rapid generation of custom vaccines by simply exchanging the antigen “plug”. The assembly pipeline involves a multi-step bioconjugation cascade: crosslinking of N-hydroxysuccinimide ester (NHS) and maleimide, third-generation copper-free click chemistry, and targeted protein ligation via the DogTag/ DogCatcher system. Our engineered DogTag scaffold (5×DT) incorporates five tandem repeats of a 23-amino-acid DogTag interspersed with flexible GGGS linkers and a C-terminal cysteine for crosslinking of *trans*-cyclooctene-polyethylene glycol (PEG_3_)-maleimide (TCO-MAL). Complementary target antigens are genetically fused to the DogCatcher domain (DC) and produced in an *E. coli* protein expression system. Initially, tetrazine-NHS (Tz-NHS) was used to introduce Tz onto the primary amines of the MV surface (Tz-MVs). The TCO-labeled 5×DT scaffold (TCO-DT) was subsequently anchored onto the Tz-MVs via an inverse electron-demand Diels–Alder click reaction (Blackman *et al*., 2008, Luu *et al*., 2024). Finally, the DC-fused recombinant antigen was conjugated to the 5×DT- anchored MVs (DT-MVs). The DT/ DC system spontaneously forms a specific, irreversible isopeptide bond under physiological conditions (Kang & Baker, 2011, Keeble *et al*., 2022). This covalent linkage prevents spontaneous antigen dissociation *in vivo*, ensuring that the target antigen and the self-adjuvanting MVs are delivered to immune cells in a highly coordinated manner.

### Screening of MV-producing hosts and evaluation of intrinsic adjuvanticity

To identify an optimal, highly productive, and immunologically potent carrier for our universal vaccine platform, we systematically screened MVs derived from various non-pathogenic Gram-positive strains, primarily in the genus *Bacillus*. The selection criteria were based on MV yield and intrinsic adjuvanticity. Initially, after a 24-h culture of the *Bacillus* species in brain-heart infusion (BHI) medium, MVs were isolated from the culture supernatant and quantified. Of the bacteria we tested, *B. subtilis* 168, *B. subtilis* natto BEST195, *B. subtilis* natto Miyagino, *B. licheniformis* JCM2505, and *B. pumilus* JCM2508 yielded MV levels suitable for scalable vaccine manufacturing (Figure S1). To profile their immunogenic potential as vaccine carriers, BALB/c mice were intranasally administered an admixture of these candidate MVs and ovalbumin (OVA). Subsequent enzyme-linked immunosorbent assay (ELISA)-based analysis revealed that these *Bacillus* MVs profoundly activated the mice’s immune system. Remarkably, *B. subtilis* 168, *B. subtilis* natto BEST195, and *B. subtilis* natto Miyagino exhibited potent adjuvanticity, substantially promoting the production of systemic immunoglobulin (Ig)G (in serum and bronchoalveolar lavage fluid [BALF]) and mucosal secretory IgA (sIgA) (in saliva and nasal lavage fluid [NLF]) (Figure S2). Given their remarkable capacity to drive sIgA production, MVs from these *B. subtilis* strains were deemed as highly promising carriers for intranasal vaccines. The magnitude of the induction was comparable to that elicited by MVs from a flagella-deficient clone of the probiotic *E. coli* Nissle 1917 (Hirayama & Nakao, 2020), a well-documented benchmark for highly immunogenic MV carriers (Uchiyama *et al*., 2024), whereas OVA alone or an admixture of OVA and *Porphyromonas gingivalis* OMVs failed to facilitate OVA-specific antibody production. Despite the excellent immunological profiles of the *B. subtilis* natto strains (BEST195 and Miyagino), their supernatants were prohibitively viscous due to the presence of polyglutamic acids, hindering downstream large-scale purification. Consequently, *B. subtilis* 168, a model Gram-positive bacterium renowned for its high genetic tractability, was selected as the MV-producing host in this study. We found that the *B. subtilis* MV fraction obtained from the culture supernatant contained flagella and a dark-red pigment (pulcherrimin). To further refine the purity of the MVs, we engineered chromosomal deletions of the *hag* and *cypX−yvmC* genes, which encode flagellin and the biosynthesis enzymes for pulcherrimin, respectively. We designated this modified strain as *<u>B</u>. subtilis* <u>d</u>eletional mutant for <u>M</u>V production 1 (BDM1). These genetic modifications successfully eliminated flagellar components and pulcherrimin from the MV fraction, yielding a highly purified and standardized vesicle population (Figure S3). The MVs from *B. subtilis* BDM1 were used for all subsequent vaccine formulations in this study.

### Trial manufacture of an MV vaccine using a model Antigen

Prior to the evaluation of pathogen-specific target antigens, we used mScarlet-I fluorescent protein (mSc) as a model antigen to optimize bioconjugation parameters and *in vivo* vaccination regimens. To evaluate bioconjugation between mSc fused to DogCatcher (mSc-DC) and the 5×DogTag scaffold (5×DT), purified mSc-DC and 5×DT were incubated overnight at room temperature in phosphate-buffered saline (PBS) at varying molar ratios, and the reaction products were analyzed via sodium dodecyl sulfate– polyacrylamide gel electrophoresis (SDS-PAGE). As the mSc-DC ratio increased, a progressive ladder of five distinct high-molecular-weight bands emerged, indicating the formation of polymerized mSc-DC and 5×DT conjugates (Figure 2A). Their molecular weights did not match those deduced from the protein size markers because the conjugates were branched owing to isopeptide bonds between DT and DC. This *in vitro* assay confirmed the successful polymerization of mSc-DC onto 5×DT. Subsequently, we verified the click chemistry-mediated anchoring of the *trans*-cyclooctene-labeled 5×DT (TCO-DT) onto the tetrazine-labeled *B. subtilis* MVs (Tz-MVs). After the incubation of TCO-DT with the Tz-MVs, Western blot analysis showed a pronounced high-molecular- weight shift for TCO-DT (Figure 2B). The mobility shift depended on the TCO and Tz labeling, as the control mixtures with unlabeled MVs showed no such interaction, definitively proving the stable covalent anchoring of TCO-DT. The broad signal of the conjugated TCO-DT strongly suggests its widespread anchoring across a heterogeneous population of the MV surface molecules (Figure 2B, lanes 4 and 5). For the final vaccine formulation, 5×DT-anchored MVs (DT-MVs) were reacted with the purified mSc-DC at room temperature in PBS. Western blot analysis using an anti-mSc antibody showed a marked upward shift of the mSc-DC band, indicating the successful anchoring of the antigen onto the DT-MVs (Figure 2C, lanes 4 and 5). Densitometry of the free mSc-DC signal in the mixture with DT-MVs indicated that more than 70% of mSc-DC in the mixture were anchored to DT-MVs, although the mSc-DC multimerization on the 5×DT scaffold did not universally result in complete 5-mer saturation, as inferred from the molecular weights of the shifted bands. Morphological assessment via field-emission scanning electron microscopy (FE-SEM) revealed that the formulated mSc-MV vaccine had a roughened surface architecture with small protrusions, in contrast to the unlabeled MVs (Figure 3A). Hypothesizing that these small protrusions corresponded to the conjugated mSc-DC (and the underlying 5×DT), we performed immunogold labeling by using an anti-mSc primary antibody and a gold-conjugated secondary antibody. Immunogold electron microscopy detected the gold particles on the surface protrusions (Figure 3B, mSc-MV). The additional textural roughness observed on the immunolabeled mSc-MV further corroborated the dense binding of the antigen–antibody complexes. Taken together, these imaging data demonstrate the successful anchoring of mSc-DC onto the MV carrier.

**Figure 2.**
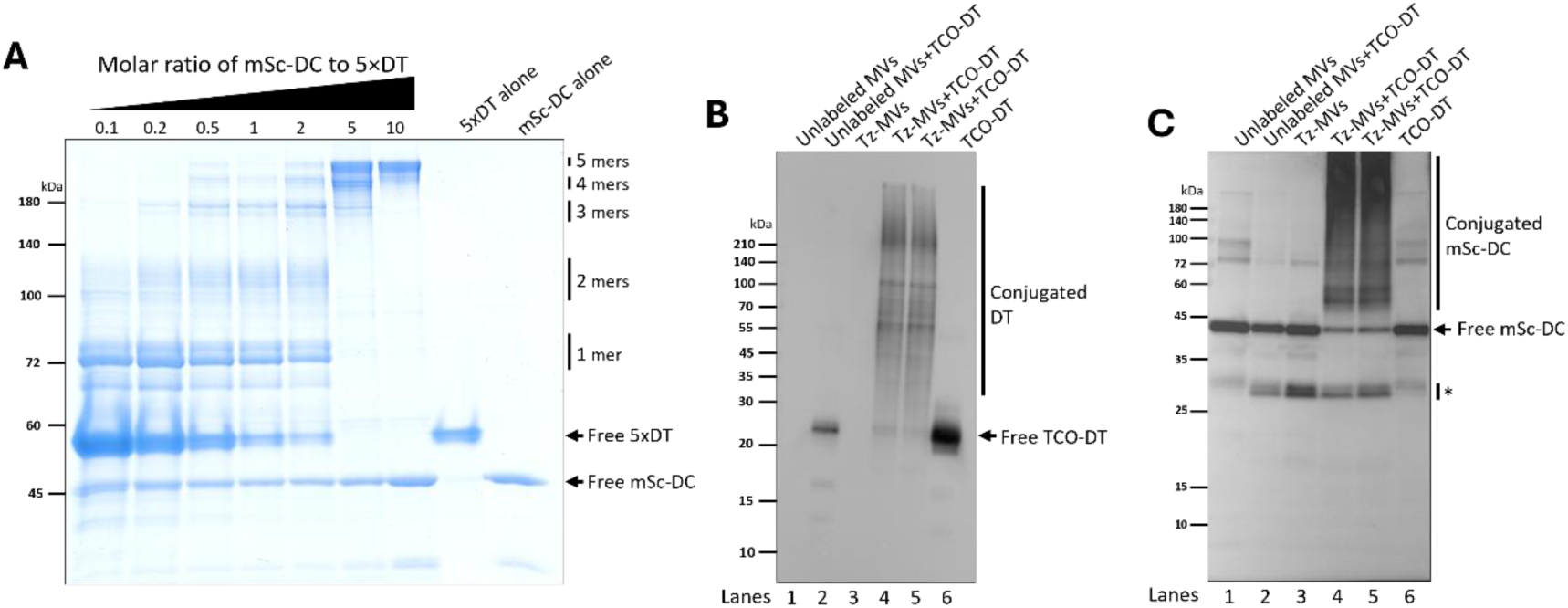
Development of the MV vaccine platform. (**A**) Multimerization of mScarlet fused to DogCatcher (mSc-DC) on the 5×DogTag scaffold (5×DT). Recombinant mSc- DC and 5×DT were incubated at room temperature in PBS at the indicated ratios, decreasing the concentrations of 5×DT. The reaction products were analyzed via SDS-PAGE followed by Coomassie brilliant blue (CBB) staining. The deduced valency of the mSc-DC multimers is indicated as -mers. Unlabeled 5×DT tends to migrate slowly due to disulfide bond formation. (**B**) Anchoring of 5×DT onto Tz-MVs. TCO-DT and Tz-MVs were incubated at room temperature in PBS. Anchoring of 5×DT onto the Tz-MVs was verified via Western blotting by using an anti-His_6_ antibody. Lane 1, *B. subtilis* MVs without Tz label; lane 2, a mixture of *B. subtilis* MV without Tz label and TCO-DT; lane 3, Tz-MVs alone; lanes 4 and 5, a mixture of Tz-MVs and TCO-DT (two independent lots prepared in parallel); lane 6, TCO-DT alone. (**C**) Anchoring of mSc-DC onto DT- MVs. mSc-DC was incubated with DT-MVs in PBS. Anchored mSc-DC was detected using an anti-RFP antibody. The arrow and asterisk indicate free (unbound) and truncated mSc-DC, respectively.

**Figure 3.**
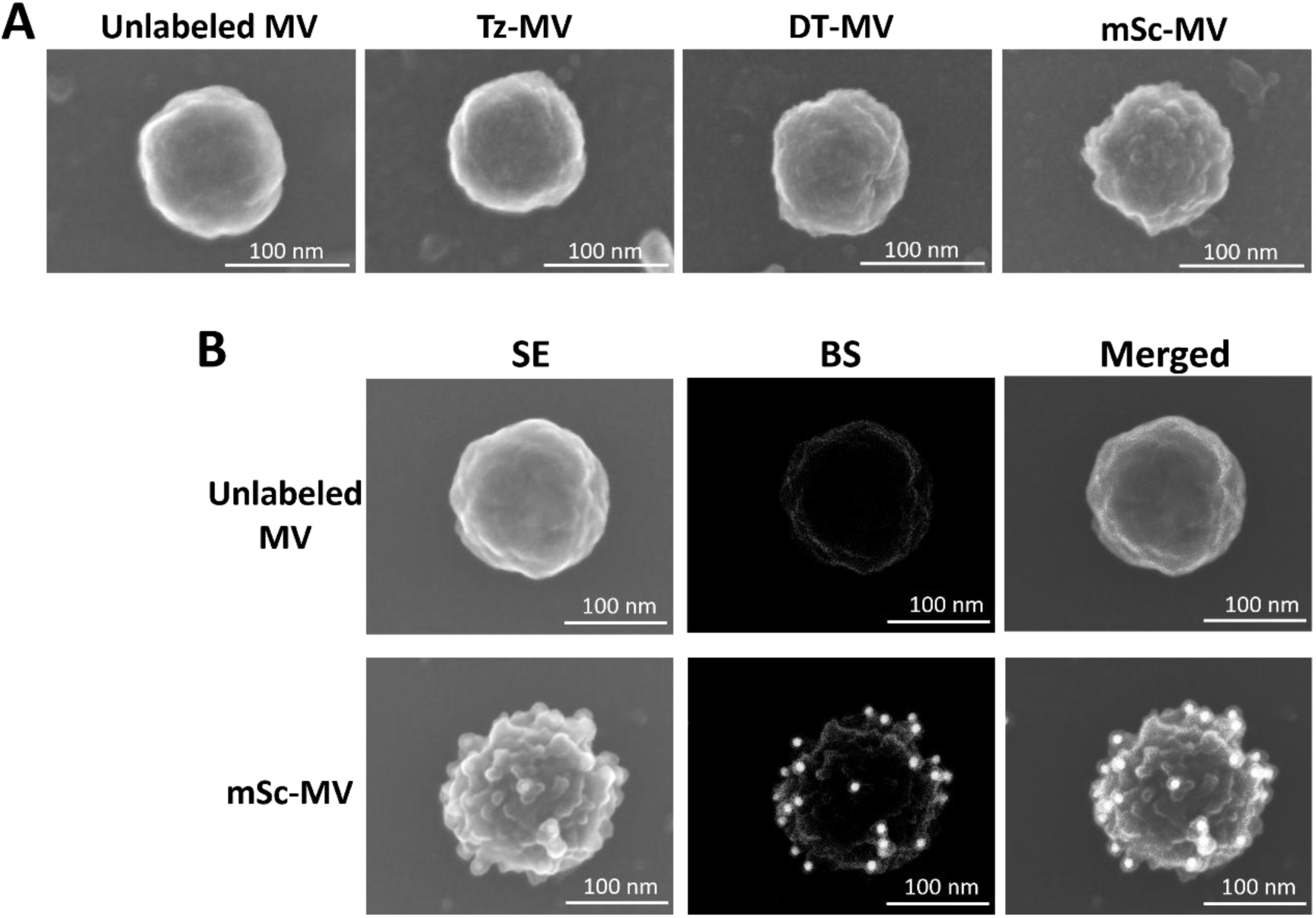
FE-SEM images of mSc-MVs. **(A**) *B. subtilis* MV samples from each step of the vaccine production process were observed using field emission scanning electron microscopy (FE-SEM). Scale bar, 100 nm. (**B**) Immunodetection of mSc-DC on mSc- MVs. Unlabeled MVs and mSc-MVs were subjected to FE-SEM imaging after immunoreaction with the anti-RFP primary antibody and a gold particle-conjugated secondary antibody. SE, secondary electron image; BE, backscattered electron image; Merged, merged image of SE and BE. White dots indicate the gold particles conjugated to the secondary antibody. Scale bar, 100 nm.

### Immunogenicity profiling of mSc-MVs

Using the formulated mSc-MV, we characterized the immunological profile of the MV vaccine platform in a murine model. To evaluate the dose-dependency of the induced immune response, 6-week-old female BALB/c mice were intranasally administered mSc- MVs. The immunization regimen consisted of a primary dose followed by two booster doses at 2-week intervals. To assess the immunological significance of the covalent linkage between the mSc-DC antigen and the MV carrier, a control group was immunized with a simple admixture of mSc-DC and unlabeled MVs by using the same regimen. In addition, another control group received an admixture of mSc-DC and aluminum hydroxide gel (Alum), one of the most commonly used adjuvants in humans. Post-vaccination analyses of serum, saliva, bronchoalveolar lavage fluid (BALF), and nasal lavage fluid (NLF) demonstrated that intranasal administration of mSc-MVs elicited high levels of antigen-specific IgG (in the serum and BALF) and sIgA (in the NLF and saliva), confirming the successful induction of both systemic and mucosal immunity (Figure 4A and 4B). Statistical evaluation revealed a dose-dependent increase in both IgG and sIgA production, peaking at the 5 μg dose. By contrast, equivalent doses of mSc-DC alone or mSc-DC adjuvanted with Alum failed to elicit comparable immune responses. Furthermore, mice immunized with the admixture of mSc-DC and unlabeled MVs exhibited negligible mSc-specific antibody titers, highlighting the importance of the covalent linkage between the antigen and the MV carrier for effective co-delivery. Unexpectedly, while the admixture of mSc-DC and unlabeled MVs elicited substantial background reactivity against the MV carrier backbone, this off-target response was suppressed in mice immunized with the conjugated mSc-MV (Figure 4C and 4D).

**Figure 4.**
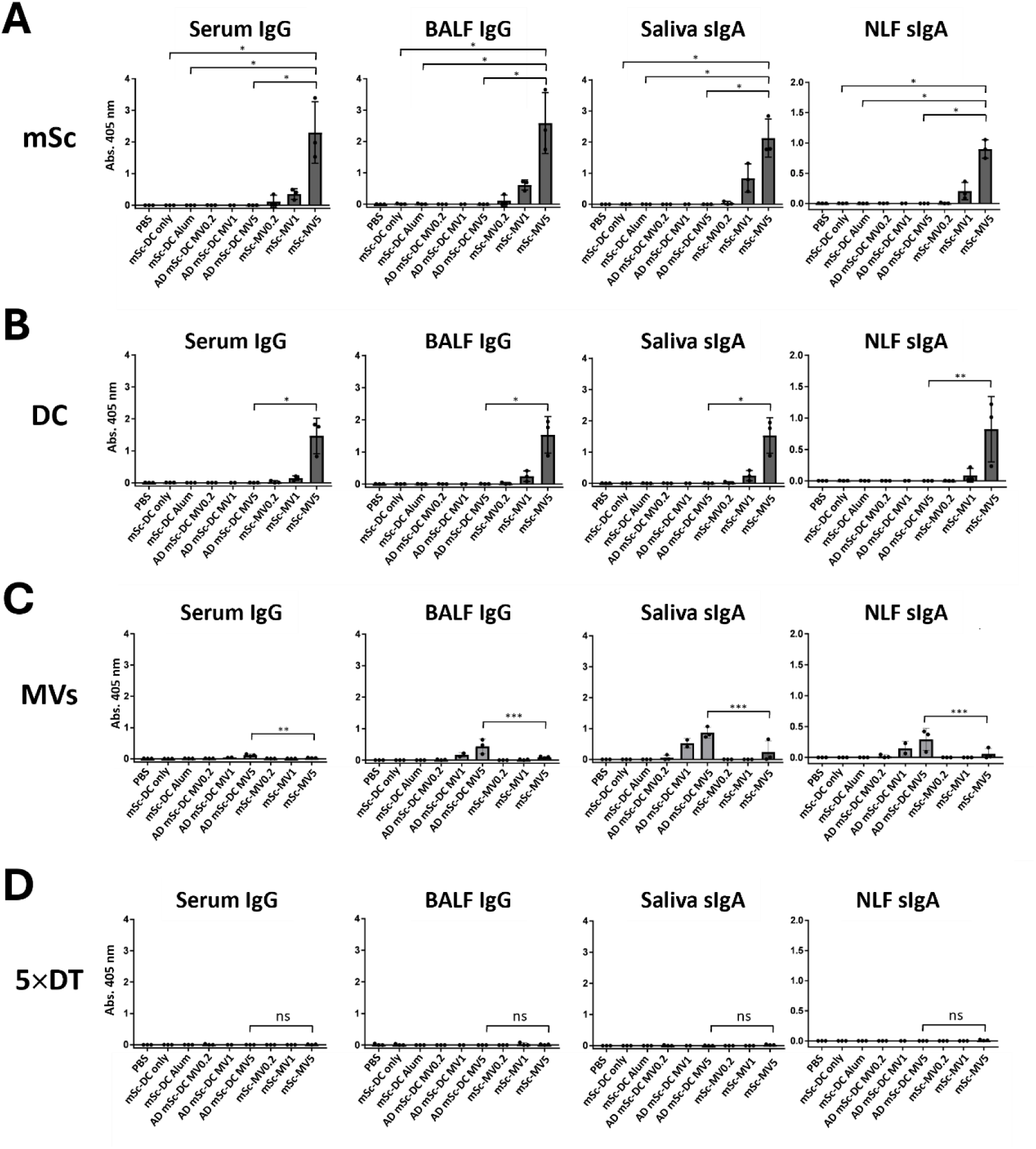
*In vivo* immunogenicity of mSc-MVs. Mice were intranasally administered mSc-MVs three times at 2-week intervals at the following doses: 0.2, 1, and 5 μg (mSc- MV0.2, mSc-MV1, and mSc-MV5). For comparison, other groups of mice were intranasally administered PBS, 5 μg of mSc-DC only, 5 μg of mSc-DC formulated with aluminum hydrogel adjuvant (100 μg/ dose; mSc-DC Alum), and simple admixtures of mSc-DC and unlabeled MVs (AD mSc-DC MV0.2, 1, and 5). One week after the third shot, an ELISA was performed to evaluate the production of IgG (in serum and BALF) and sIgA (in saliva and NLF) specific to mSc (**A**), DC (**B**), the MV backbone (**C**), and 5×DT (**D**). *n* = 3; ns, no significance; \**P* < 0.0001; \*\**P* < 0.001; \*\*\**P* < 0.03.

Multimerization of antigens can enhance their immunogenicity by facilitating the cross-linking of B cell receptors (Bachmann & Jennings, 2010). To rule out the possibility that the enhanced immunogenicity of mSc-MVs was attributed to the multimerized mSc- DC rather than the covalent binding of mSc-DC to DT-MVs, we immunized mice with an admixture of the multimerized mSc-DC bound to 5×DT and unlabeled MVs. Post-vaccination analysis revealed that the multimeric mSc-DC unanchored to the MVs failed to induce mSc-specific antibody production, underscoring the importance of the covalent bond between the antigen and the MVs (Figure S4). We further investigated the structural influence of the DT repeat length (1, 5, and 10 repeats) on immunogenicity. Consistent with our finding that covalent anchoring of the antigen to the MVs, rather than multimerization, is the primary driver of immune activation, even the single-DT construct induced an immune response (Figure S5A, mSc-DC+1×DT-MV). Nevertheless, our data indicated that an architecture comprising 5 repeats of DT maximized induction of sIgA (in the saliva and NLF) and IgG (in the BALF), although the differences between the 1- and 5-repeat constructs did not reach statistical significance. By contrast, extending DT to 10 repeats significantly attenuated the immune response, likely due to steric hindrance and subsequent MV aggregation (Figure S5B). Therefore, a standardized protocol in which we used the MV carrier with 5×DT, a 5 μg dose (5 μg of antigens and 5 μg of DT- MVs), and the intranasal route was firmly established for all subsequent therapeutic evaluations.

### Targeted application: formulation and immunogenicity of the Y. pestis mCaf1 vaccine

To validate the clinical applicability of our MV vaccine platform, we formulated a vaccine targeting *Y. pestis*. Despite extensive historical efforts, no licensed plague vaccine is currently available for public use (Inglesby *et al*., 2000). Rational vaccine design against *Y. pestis* often uses the protective Caf1 capsular polymeric protein (F1 antigen) (Williamson, 2001). We engineered a recombinant Caf1 protein fused to DC. To prevent spontaneous polymerization and to maintain a monomeric state, mCaf1 was used as described previously (Chalton *et al*., 2006). mCaf1-DC was covalently anchored to the DT-MV carrier to generate mCaf1-MVs. Immunological profiling following intranasal administration revealed a stark contrast in immunogenicity: mCaf1-MVs induced robust production of mCaf1-specific IgG and sIgA, whereas mCaf1 alone and mCaf1 adjuvanted with Alum did not elicit immune responses (Figures 5A and S6), highlighting the potent adjuvanticity of the MV carrier in our system. Consequently, we focused on the highly immunogenic mCaf1-MV vaccine for detailed analysis. IgG subclass analysis demonstrated that mCaf1-MVs successfully orchestrated a balanced immune response, evidenced by the simultaneous induction of Th1-associated (IgG2a and IgG2b) - and Th2-associated (IgG1) isotypes (Figure 5B). A balanced Th1/Th2 response is highly desirable for the clearance of intracellular and extracellular pathogens, promoting both opsonophagocytosis and cytotoxic T-cell activation (Kidd, 2003). Additionally, the upregulation of IgG3 production was marginal in mice immunized with mCaf1-MVs. Importantly, the formulation exhibited an excellent safety profile, with IgE levels remaining undetectable after the final immunizations (Figure 5C). The absence of IgE production confirms the non-allergenic profile of the MV platform, distinguishing it from certain chemical adjuvants that can skew immunity toward reactogenic Th2 pathways.

**Figure 5.**
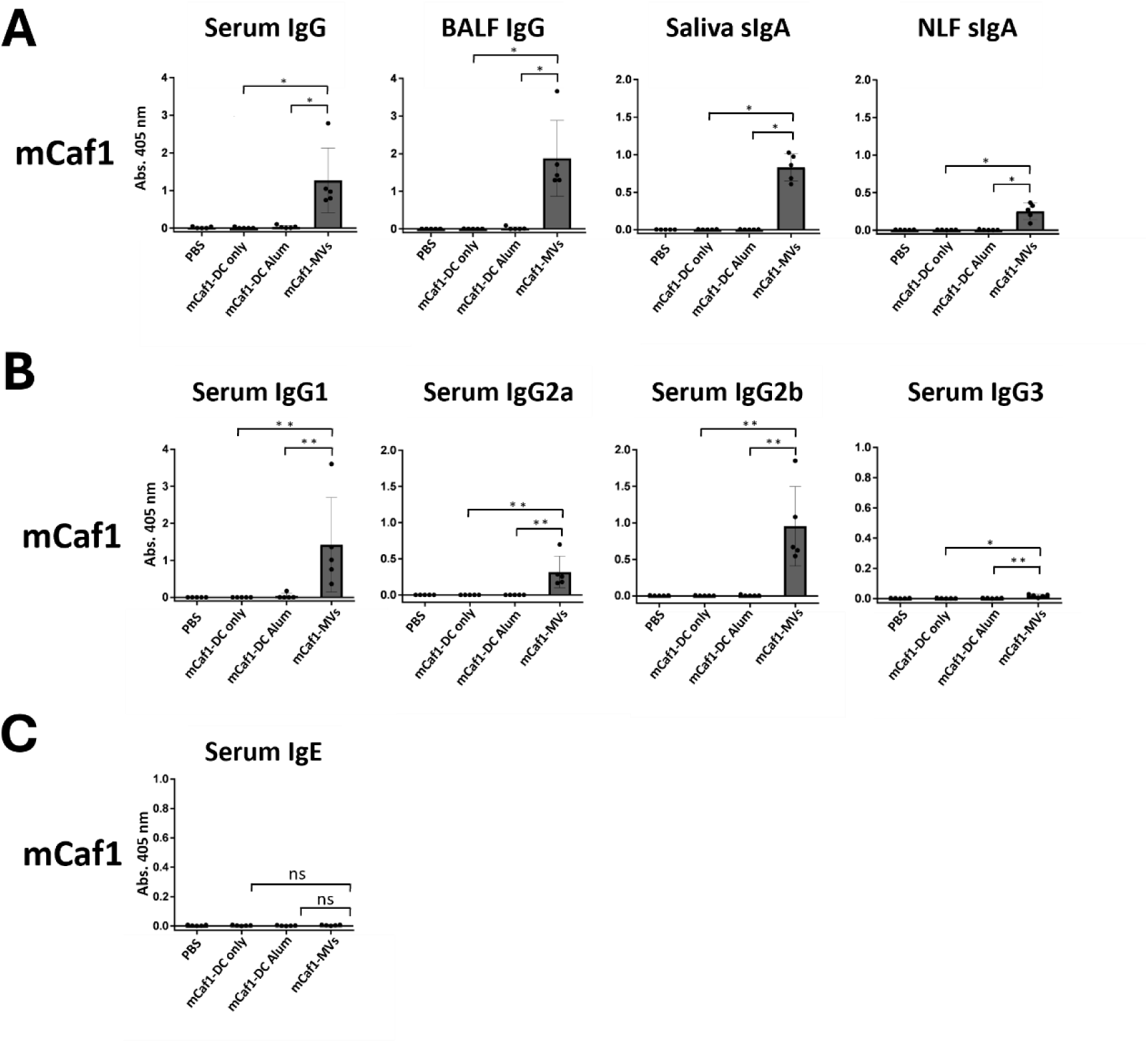
Immunogenicity of the *Y. pestis* vaccine. **(A)** Mice were intranasally immunized with the mCaf1-MV vaccine. Production of IgG (in serum and BALF) and sIgA (in saliva and NLF) specific to mCaf1 was evaluated via ELISA. (**B**) The Th1/Th2 balance was evaluated by quantifying IgG subclasses specific to mCaf1 in sera from mice immunized three times with PBS, mCaf1-DC, mCaf1 adjuvanted by Alum, or mCaf1- MVs. *n* = 5. (**C**) Serum IgE levels in mice immunized three times with PBS, mCaf1-DC, mCaf1 adjuvanted by Alum, and mCaf1-MVs were measured via ELISA. *n* = 5; ns, no significance; \**P* < 0.0001; \*\**P* < 0.01; \*\*\**P* < 0.05.

### Protection efficacy against lethal Y. pestis challenge

The definitive validation of the protective efficacy of the mCaf1-MV vaccine was conducted using a highly stringent, lethal *in vivo* challenge model. Mice were immunized as shown in Figure 6A, alongside multiple control groups (PBS, mCaf1-DC alone, and mCaf1-DC adjuvanted with Alum). After confirmation of mCaf1-specific IgG production (Figure S7), the immunized mice were intraperitoneally challenged with the highly virulent *Y. pestis* strain Alexander at 10 and 100 times the median lethal dose (LD_50_). Survival kinetics clearly demonstrated the marked superiority of mCaf1-MVs (Figure 6B and 6C). As anticipated, all mice in the control and Alum-adjuvanted cohorts rapidly succumbed to fulminant septicemia within 4 days of infection. By contrast, mCaf1-MV- immunized mice exhibited significant survival advantages, achieving 100% protection against the 10 × LD_50_ challenge and an 80% survival rate against the 100 × LD_50_ challenge. The complete failure of the Alum-adjuvanted mCaf1-DC compared with the mCaf1-MV formulation demonstrates that this modular MV platform, administered intranasally, provides an unparalleled defense against highly lethal bacterial pathogens.

**Figure 6.**
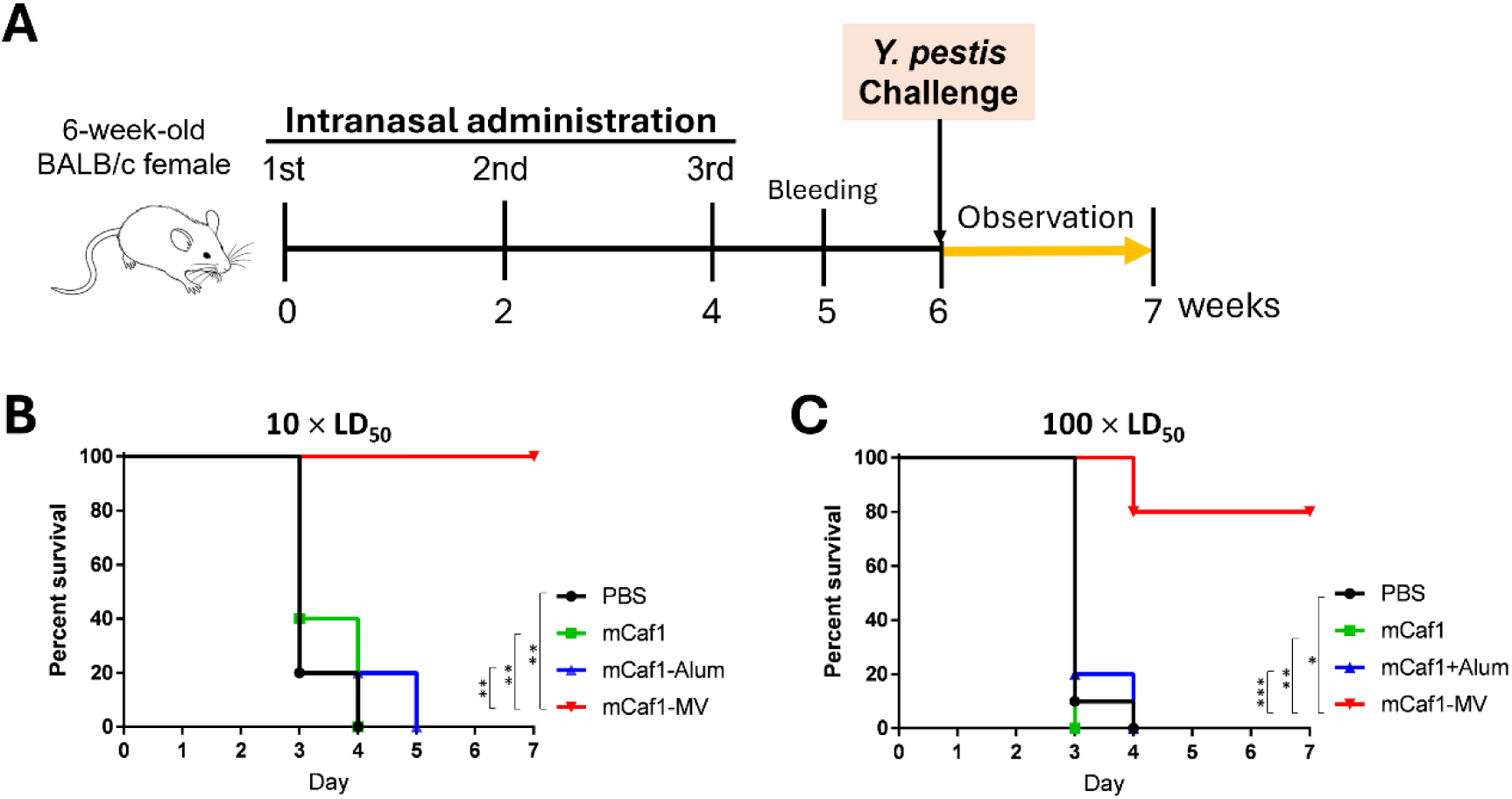
Survival assay. Mice were immunized with mCaf1-MVs three times with 2- week intervals. The immunized mice were challenged intraperitoneally with *Y. pestis* Alexander at doses of 10× (**A**) and 100× (**B**) LD_50_. Symbols: circle, PBS; triangle, mCaf1- DC, square, mCaf1-DC adjuvanted by Alum; diamond, mCaf1-MVs. *n* = 5; \**P* < 0.001; \*\**P* < 0.0025; \*\*\**P* < 0.005.

## Discussion

The rapid development of a safe, highly efficacious, and adaptable vaccine platform is a critical global health priority, particularly in the combat against highly lethal pathogens. In this study, we successfully engineered a modular, universal vaccine platform by using MVs derived from non-pathogenic *B. subtilis*. We established a “plug-and-display” covalent conjugation system and an *in vivo* vaccination regimen by using mSc-MVs. Furthermore, we demonstrated that the mCaf1-MV vaccine elicits robust mucosal and systemic immunity, providing high-level protection against a lethal plague challenge in mice. This vaccine not only provides a novel countermeasure against *Y. pestis* but also offers a versatile technological foundation potentially applicable to a broad spectrum of infectious diseases.

Historically, the exploration of bacterial vesicles in vaccinology was focused on MVs derived from Gram-negative bacteria. Although they are highly immunogenic, their clinical application is often hindered by the inherent toxicity of LPSs (Ellis & Kuehn, 2010). Additionally, MVs are associated with the export and delivery of toxins such as cytolysin A and Shiga toxin in *E. coli* and are therefore called “type zero secretion systems” (Guerrero-Mandujano *et al*., 2017). To mitigate these severe reactogenic risks, chemical and genetic detoxification processes are typically required (Gerritzen *et al*., 2017). By contrast, in this study, we pivoted toward MVs from *B. subtilis*, a Gram-positive, nonpathogenic, and GRAS bacterium, to circumvent this problem. While lacking an outer membrane, Gram-positive bacteria secrete cytoplasmic MVs, which are rich in immunostimulatory lipoproteins and peptidoglycan fragments, serving as potent TLR agonists without the endotoxic risks associated with LPS (Brown *et al*., 2015, Briaud & Carroll, 2020). Although they lack highly immunogenic LPSs, *B. subtilis* MVs exhibited intrinsic adjuvanticity, comparable to that of *E. coli* (Figure S2). Moreover, intranasal administration elicited a mucosal immune response and sIgA production as well as systemic immunity. Thus, it has the potential to create a robust frontline barrier that neutralizes pathogens at their primary entry point, providing a major protective advantage, particularly against respiratory infections. By engineering the genetically tractable *B. subtilis* 168 to lack flagellin and pulcherrimin, we secured a highly pure vesicle population.

A defining innovation of our platform is its modular “plug-and-display” architecture, using the covalent, directional anchoring of target antigens via click chemistry and the DT/ DC protein ligation system. Remarkably, the final vaccine formulation simply consists of mixing the DT-MVs with the DC-fused target antigen in PBS to achieve spontaneous covalent linkage. By contrast, conventional strategies for the development of MV-based vaccines with heterogeneous antigens involve the simple admixture of MVs with antigens or the genetic fusion of antigens to outer membrane proteins (Gerritzen *et al*., 2017, Garling *et al*., 2025). In the admixture strategy, the efficacy of the vaccines heavily relies on the intrinsic immunogenicity of the antigens. In the context of a universal vaccine platform, antigen-dependent immunogenicity is highly undesirable; rather, the system must elicit consistent vaccine efficacy regardless of the target antigen. Indeed, while the MV admixtures of the MVs with OVA successfully stimulated OVA-specific antibody production (Figure S2), the covalent anchoring of the antigens to DT-MVs was required to elicit an immune response in our study (Figures 4A and S4A). Another conventional strategy is the genetic fusion of heterologous antigens to native outer membrane-anchoring proteins such as OmpA in *E. coli* (König *et al*., 2021, Tong *et al*., 2024). This genetic fusion approach, however, is fraught with limitations: the expression of large, complex antigens often imposes a severe burden on the host bacterium, leading to drastically reduced MV yields, antigen misfolding, or degradation (Chen *et al*., 2010). Our platform resolves these bottlenecks by decoupling MV production from antigen expression. Specifically, we used the *B. subtilis* MVs as the vaccine carriers and recombinant antigens produced in *E. coli*. Crucially, our approach is highly adaptive; it can be readily extended to incorporate glycosylated antigens in eukaryotic protein expression systems, which bacterial hosts are inherently incapable of synthesizing.

*Y. pestis* can cause human-to-human transmission via respiratory droplets from infected patients, posing the potential to trigger a pandemic. For pathogens such as *Y. pestis*, the induction of mucosal immunity in the respiratory tract is as critical as the systemic humoral defense (Smiley, 2008). sIgA serves as the first line of defense, neutralizing the pathogen at the mucosal epithelium before systemic dissemination can occur. Intranasal administration of the mCaf1-MV vaccine led to effective production of both systemic IgG and mucosal sIgA, sufficient to protect immunized mice from a *Y. pestis* challenge (Figures 5, 6, and S7). Although researchers have reported that intramuscular administration of mCaf1 and Alum effectively stimulated an immune response (Chalton *et al*., 2006), Alum did not enhance the immunogenicity of mCaf1 in our study (Figure 5 and S7). Likely, the Alum-mCaf1 complex is not invasive to the mucosa in the nasal cavity because Alum forms insoluble particles with antigens (Laera *et al*., 2023). These results demonstrate the efficacy and versatility of the mCaf1-MV vaccine.

Despite its efficacy, the mCaf1-MV vaccine alone may be impractical to use due to the risk of “antigenic escape” driven by spontaneous *caf1* mutations in *Y. pestis* (Friedlander *et al*., 1995). To overcome this vulnerability, an effective plague vaccine requires the co-delivery of alternative antigens. Beyond Caf1, other *Y. pestis* antigens have been suggested for use in vaccines, including LcrV, YscF, and Yop proteins (type III secretion system components and effector proteins) (Williamson *et al*., 2024). As previous studies have shown that a genetic fusion of *caf1* and *lcrV* effectively elicits immune responses in mice (Heath *et al*., 1998, Goodin *et al*., 2007), anchoring a single LcrV- mCaf1-DC fusion antigen to DT-MVs is a highly promising strategy for the development of a multivalent vaccine. Combining mCaf1 with those antigens may constitute a powerful strategy to create broadly protective, multivalent *Y. pestis* vaccines. Our highly modular MV vaccine platform is ideally suited to readily incorporate these diverse antigens, thereby substantially accelerating the development of next-generation vaccines against plague and other pathogens.

## Materials and Methods

### Bacterial strains and culture conditions

Bacterial strains and plasmids are listed in Table S1. *Bacillus* strains were cultured at their optimal temperature in Luria–Bertani (LB) medium (#244620; Becton Dickinson and Company [BD], MD, USA) for routine culture or in Brain-Heart infusion medium (BHI; #237500; BD) for MV production. *B. subtilis*, *B. subtilis* natto, *B. licheniformis*, and *B. velezensis* were cultured at 37 °C, and *B. megaterium*, *B. pumilus*, *B. weihenstephanensis*, and *B. mojavensis* were cultured at 30 °C. *E. coli* strains JM109 (#9052; TAKARA, Shiga, Japan) and BL21(DE3) SHuffle T7 Express *lysY* (#3030J; New England BioLabs [NEB], MA, USA) were used for plasmid cloning and protein expression, respectively. *E. coli* strains harboring pET22b(+)-derived vectors (#69744; Merck Millipore, MW, USA) or its derivative plasmids were cultured in LB medium containing 100 μg/ml ampicillin. *Y. pestis* strain Alexander (Fukui *et al*., 1957) was cultured on BHI-agar plates at 28 °C for recovery from frozen stocks or at 37 °C to induce virulence expression.

### Isolation of MVs

*Bacillus* strains were precultured overnight at their optimal temperature, as described above, in 10 ml of LB medium. One milliliter of the precultures was inoculated into 100 ml of fresh BHI medium and cultured at their optimal temperature for 24 h with shaking. These cultures were centrifuged at 9,000× *g* and 4 °C for 10 min. The culture supernatants were filtered through a 0.45-μm polyvinylidene fluoride (PVDF) filter (#S2HVU01RE; Merck Millipore) and subsequently ultracentrifuged at 150,000× *g* and 4 °C for 1 h. The MV pellets were washed with PBS, again ultracentrifuged at 150,000× *g* and 4 °C for 1 h, and resuspended in PBS. Protein contents of the MV samples were measured using a bicinchoninic acid (BCA) quantification kit (#T9300A; TAKARA) with 0.1−2.0 mg/ml bovine serum albumin (BSA) standards. Lipid contents of the MV samples were quantified using FM1-43 dye (#T3163; Thermo Fisher Scientific, MA, USA), as described previously (Abe *et al*., 2021). In brief, 5 μl of each MV sample was mixed with 100 μl of 1.25 μg/ml FM1-43 dye. FM1-43 fluorescence (excitation/emission, 472/580 nm) was measured using a Cytation 5 microplate reader (Biotek, VT, USA).

### Construction of *B. subtilis* strain BDM1 for MV production

A *B. subtilis* mutant strain lacking *hag* and *yvmC−cypX* (designated as BDM1, *<u>B</u>. subtilis* <u>d</u>eletional mutant for <u>M</u>V production 1) was created to use for the MV production. Spectinomycin (*spec*)- and erythromycin (*ermC*) resistance gene cassettes were created to delete the *hag* and *yvmC−cypX* genes of *B. subtilis* 168. The 5′- and 3′-franking regions of *hag* and *yvmC−cypX* were amplified from the *B. subtilis* genome by using the primer sets, PKA-837/739 (for 5′-*hag*, listed in Table S2), PKA-313/314 (3′-*hag*), PKA-070/071 (5′- *yvmC*), and PKA-072/073 (3′- *cypX*), respectively. The *spec* and *ermC* genes were obtained via PCR of the genomic DNA from *B. subtilis* Δ*lytCDEF* strain (Abe *et al*., 2021) by using the PKA-254/255 and PKA-009/010 primer sets. The *spec* cassette for *hag* deletion and *ermC* cassette for *yvmC−cypX* deletion were generated by overlap-extension PCR (OE-PCR) with PKA-837/314 and PKA-070/073, respectively. The *spec* and *ermC* cassettes were introduced into *B. subtilis* 168 via natural competence and selected by 100 μg/ml spectinomycin and 1 μg/ml erythromycin.

### Construction of expression plasmid vectors

To add a C-terminal six-histidine tag (His_6_) to the recombinant proteins, the pET22b(+) plasmid was used in this study. A DNA fragment encoding five repeats of DogTag (5×DT) interspersed with a (GGGS)_2_ linker sequence was synthesized by Eurofins Genomics (Tokyo, Japan) and used as a PCR template. For pFLAG-5×DT, the 5×DT DNA fragment was amplified with a primer set PKB-334/371, inserted into the *Nde*I/*Xho*I site of pET22b(+) vector via Gibson assembly (NEBuilder, #E2621; NEB), and cloned into *E. coli* JM109. The resulting plasmid was linearized via inverse PCR with PKB-479/480 to add the 5′-terminal 3×FLAG tag (DYKDHDGDYKDHDIDYKDDDDK)-coding sequence and self-ligated with T4 polynucleotide kinase (#312-01551, Nippon Gene, Tokyo, Japan) and T4 DNA ligase (DNA Ligation Kit Mighty Mix, #6023; TAKARA). For pFLAG-1×DT, an inverse PCR was performed using pFLAG-5×DT and PKB- 478/481, followed by self-ligation with T4 ligase. For pFLAG-10×DT, a 5×DT DNA fragment was amplified with PKB-493/492, digested with *Bam*HI, and inserted into the *Bam*HI site of pFLAG-5×DT. A DNA fragment encoding (GGGS)_2_-linker−DogCatcher (DC), synthesized by Eurofins Genomics, was amplified with PKB-563/345, inserted into the *Nde*I/*Xho*I site of pET22b(+) vector via Gibson assembly, and cloned into *E. coli* JM109. For pET-mSc, the PKB-352/137 primer set was used to amplify the mScarlet-I gene. The PCR product was inserted into the *Nde*I/*Xho*I site of the pET22b(+) vector via Gibson assembly and cloned into *E. coli* JM109. For an mScarlet-I (mSc)−DC fusion gene, mSc and (GGGS)_2_-linker−DC were amplified with PKB-352/148 and PKB- 134/345, respectively. These DNA fragments were combined via OE-PCR, inserted into the *Nde*I/*Xho*I site of the pET22(b)+ vector, and cloned into *E. coli* JM109. For the production of a circularly permutated form of *caf1* (encoding mCaf1), DNA fragments were generated via PCR from the *Y. pestis* genome by using the PKB-310/311, PKB- 312/313, and PKB-314/315 primer sets. These fragments were combined and amplified via OE-PCR with the PKB-310/315 primer set and inserted into the *Nco*I/*Xho*I site of the pET22b(+) vector. For pET-mCaf1-DC, *mcaf1* was amplified with PKB-310/345, and pET-mSc-DC was linearized via inverse PCR with PKB-134/092. The resulting DNA fragments were combined via Gibson assembly and cloned into *E. coli* JM109.

### Protein purification

*E. coli* BL21(DE3) SHuffle T7 Express *lysY* and strains harboring the expression plasmids were cultured in LB medium containing 100 μg/ml ampicillin at 30 °C, with shaking. Protein expression was induced by adding 0.5 mM isopropyl-β-D-1- thiogalactopyranoside (IPTG) at the exponential phase of cell growth, and the *E. coli* cells were cultured overnight at 22 °C. All of the proteins used in this study were His6-tagged; therefore, nickel-nitrilotriacetic acid (Ni-NTA) affinity chromatography was used for purification. For the purification of 5×DT, mSc, DC, and mSc-DC, which show cytoplasmic expression, the *E. coli* cells were disrupted using the cell lysis reagent, FastBreak (#V8571; Promega, WI, USA) in a solution containing 50 mM sodium phosphate (pH 8.0) and 0.3 M sodium chloride. The His_6_-tagged recombinant proteins were affinity-purified with a Ni-NTA column (HisTrap FF crude, #11000458; Cytiva, MA, USA). After the column was washed with a buffer containing 50 mM sodium phosphate (pH 8.0), 1 M sodium chloride, and 0.1% triton X-100, the recombinant proteins were eluted with a buffer containing 50 mM sodium phosphate (pH 8.0), 0.3 M sodium chloride, and 500 mM imidazole. For mCaf1, mCaf1-DC, which show periplasmic expression, the *E. coli* cells after the protein expression induction were resuspended in a hypertonic solution containing 50 mM Tris-HCl (pH 8.0) and 20% sucrose for 10 min on ice and centrifuged briefly. The cell pellet was resuspended in water and kept on ice for 10 min. After centrifugation, the supernatants were collected, and sodium phosphate buffer (pH 8.0) and sodium chloride were subsequently added to final concentrations of 50 mM and 0.3 M, respectively. The His_6_-tagged proteins were purified with HisTrap FF crude, as described above.

### MV vaccine production

One microgram of *B. subtilis* MVs was labeled with 1 mM tetrazine-NHS ester (Tz-NHS, #T4125; TCI, Tokyo, Japan) in 1 ml of PBS for 3 h at room temperature. After the reaction, excess Tz-NHS was removed by ultracentrifugation, followed by washing with PBS. The 5×DT protein (1 mg) was labeled with 2 mM *trans*-cyclooctene-polyethylene glycol (PEG_3_)-maleimide (TCO- MAL; #T3948, TCI) in 1 ml of PBS containing 1 mM Tris-(2- Carboxyethyl)phosphine (TCEP; T2556: Thermo Fisher Scientific) for 3 h at room temperature. To remove excess TCO-MAL, TCO-DT was purified with a Ni-NTA column. Tz-MVs and TCO-DT were reacted at the equivalent weight in PBS for 1 h at room temperature. After the reaction, TCO- DT unbound to Tz-MVs was removed via ultracentrifugation, followed by resuspension in 0.5 ml of PBS. The DT- MVs were stored at -80 °C before antigen anchoring. Equal amounts of the recombinant antigens fused with DC and DT-MVs were reacted in PBS at room temperature for 1 h and stored at 4 °C before vaccination.

### Western blotting

The recombinant proteins and the MV samples were subjected to SDS-PAGE and transferred onto PVDF membranes (Immobilon-P, #IPVH09120; Merck) by using an electroblotter (Trans-Blot Turbo Transfer System #1704150; BioRad, CA, USA). The membrane was blocked with a blocking reagent (Blocking ONE, #03953-95; Nakarai, Tokyo, Japan), reacted with anti-His6 antibody-HRP conjugate for 5×DT (Anti-His-tag mAb-HRP-DirecT; #D291-7, MBL) and anti-pan-RFP rabbit antibody for mSc-DC (#pabr1-20; Cosmo Bio Co., LTD, Tokyo, Japan). For detection of the anti-pan-RFP antibody, anti-rabbit IgG (H+L) antibody-HRP conjugate (#W4011; Promega) was used at a dilution ratio of 1:20,000. Chemiluminescence was generated using a luminescent HRP substrate (Western BLoT Hyper HRP Substrate, #T7103A; TAKARA) and detected with a cooled charged-coupled camera (Fusion Solo 5; Vilber Bio Imaging, Marne-la- Vallée, France).

### Field emission scanning electron microscopy (FE-SEM) imaging

For FE-SEM imaging, *B. subtilis* MVs were mounted on poly-L-lysine-coated coverslips immobilized in four-well multidishes (#176740; Thermo Fisher Scientific) and incubated at room temperature for 200 min to allow attachment. The MV samples were fixed overnight at 4 °C with 2.5% glutaraldehyde and 2% paraformaldehyde in PBS. The fixed MVs were dehydrated through a graded acetone series, dried using a CO_2_ critical-point dryer (CPD300, Leica Microsystems, Wetzlar, Germany), and coated using an osmium plasma coater (Neoc-Pro/P, Meiwafosis, Tokyo, Japan). Subsequently, the osmium-coated MV samples were imaged with an FE-SEM (Regulus 8220; Hitachi High-Tech, Tokyo, Japan) at an accelerating voltage of 5 kV. For immunostaining-SEM, the nonlabeled MVs and mSc-MVs were blocked with 2.5% skim milk in PBS, reacted with anti-pan-RFP antibody, and subsequently reacted with anti-rabbit IgG antibody–gold colloidal particle- conjugate (#AC-10-01-05; Cosmo Bio) prior to fixation.

### Animal experiments

All murine experiments were reviewed and approved by the Animal Use and Care Committee of the National Institute of Infectious Diseases in Japan (NIID; approved protocol No. 125165). All protocols were performed at the NIID in accordance with institutional guidelines and regulations. Specific pathogen-free (SPF), female BALB/c mice were purchased from Japan SLC Inc. (Shizuoka, Japan). All mice were randomly group-housed (3−5 mice per cage) with ad libitum access to water and a chow diet under a standardized light cycle (12 h on/12 h off) in the NIID animal facility.

### Immunization of mice

Six-week-old SPF female BALB/c mice were purchased from Japan SLC Inc. Intranasal administration of vaccines was performed at a total volume of 10 μl to both nostrils (5 μl each), under inhalation anesthesia with 2.5% isoflurane (Merk KGaA, Darmstadt, Germany). When an aluminum hydroxide adjuvant (Alum; # 012-24241; Fujifilm Wako, Saitama, Japan) was used instead of the MV vaccine, 100 μg/dose of Alum was mixed with the antigens and administered to the mice. Three rounds of vaccination were performed at 2-week intervals.

### Sampling of serum, saliva, NLF, and BALF

Whole blood was drawn at euthanasia via cardiac puncture by using a 26-gauge needle syringe under inhalation anesthesia with 2.5% isoflurane. Partial blood was collected from the tip of the tail vein after a cut was made with scissors. After coagulation, serum was recovered from the blood samples via brief centrifugation and stored at −30 °C. For saliva collection, mice were intraperitoneally injected with 100 μl of a secretagogue to stimulate the parasympathetic nervous system. The secretagogue was a mixture of 0.4 mg/ml isoproterenol (Merk Millipore) and 0.1 mg/ml pilocarpine (Merk Millipore) in PBS. 3−5 minutes after the injection, secreted saliva was collected using a micropipette without anesthesia. The collected saliva was frozen at −80 °C and thawed at room temperature, and stored at −30 °C. For NLF and BALF sampling, the upper head and lungs of euthanized mice were collected after whole blood sampling. NLF samples were obtained by washing the nasal cavity of the isolated upper head three times with 1 ml of PBS containing 0.1% BSA. BALF sample was obtained by washing the dissected lungs, including the trachea, three times with 2 ml of PBS containing 0.1% BSA.

### Enzyme-linked immunosorbent assay (ELISA)

ELISAs were performed to assess the amounts of antigen-specific antibodies in serum, saliva, NLF, and BALF samples, as described previously (Uchiyama *et al*., 2024). Purified recombinant antigens (mSc, mSc-DC, DC, 5×DT, and mCaf1) were coated onto an ELISA plate (125 ng/well) (#675061; Gainer Bio-One GmbH, Frankhauser, Germany). Alkaline phosphatase (AP)-labeled anti-mouse IgG Fcγ fragments were purchased from Jackson ImmunoResearch (#315-055-008; PA, USA), and used at a 1:1,000 dilution ratio. AP-labeled anti-mouse IgE, IgA, IgG1, IgG2a, IgG2b, and IgG3 were purchased from SouthernBiotech (Birmingham, AL, USA). The anti-mouse immunoglobulins were used at 1:1,000−2,000 dilution ratios. Chromogenic development via para-nitrophenyl phosphate at 405 nm (A_405_) was measured using Cytation 5. The A_405_ values were subtracted from the average A_405_ values of three wells incubated on the same plate with PBS alone (without any mouse samples).

### Animal challenge test

A frozen stock of *Y. pestis* Alexander was recovered on a BHI-agar plate at 28 °C for 48 h, then streaked onto another fresh BHI-agar plate, and incubated at 37 °C for 48 h to allow expression of its virulence factors, including Caf1. The colonies were scraped from the plate and resuspended in sterile saline to adjust the optical density of 1.0. The *Y. pestis* suspension was serially diluted in sterile saline before injection. One week after the three intranasal immunizations, the vaccinated mice were injected intraperitoneally with 100 μl of the *Y. pestis* suspension at 10 or 100 × times the median lethal dose (LD_50_; 1 LD_50_ < 10 c.f.u.) and observed for 1 week.

### Statistical analysis

Statistical analyses were performed using Prism software (version 7.05; GraphPad Software, MA, USA). For ELISA data, differences among multiple groups were evaluated using one-way analysis of variance, followed by Tukey’s multiple comparisons test. ELISA results are presented as the mean ± standard deviation. Survival data from the *Y. pestis* challenge assay were plotted using the Kaplan-Meier method and compared using the log-rank test. Statistical significance was defined as *P* < 0.05.

## Data availability

All data presented in this paper are available from the corresponding author upon request.

## Acknowledgments

This work was supported by the Japan Society for the Promotion of Science (KAKENHI; grant number 22K05400 to KA). We thank Fumiko Takashima for her technical support.

## Author contributions

KA designed this study and wrote the original draft of the manuscript. KA and TW constructed the bacterial strains and conducted the experiments. KA, HK, and KS performed the *Y. pestis* challenge tests. TW, HK, KS, RN, TY, TS, and YA reviewed and edited the manuscript.

## Statement

The authors declare that they have no competing interests.

