## Supplementary material for "A universal plug-and-display vaccine platform for mucosal and systemic immunity using *Bacillus* subtilis membrane vesicles": SI

### Supplementary Table 1.

Strains and plasmids used in this study.

| Strain or plasmid | Genotype and/or relevant features | Source or reference |
|---|---|---|
| **Strains** | | |
| *Bacillus subtilis* | | |
| 168 | *trpC2* | *Bacillus* Genetic Stock Center (U.S.A.) |
| BDM1 | *trpC2*, Δ*yvmC–cypX*::*ermC*, Δ*hag*::*spc*, derived from 168 | This study |
| PS216 | undomesticated wild-type strain | Durrett *et al*., 2013 |
| NCIB 3610 | undomesticated wild-type strain | National Collection of Industrial Bacteria (U.K.) |
| RO-NN-1 | undomesticated wild-type strain | Earl *et al*., 2012 |
| Miyagino | *bio*$^{-}$, natto starter strain | Nishito *et al*., 2010 |
| BEST195 | *bio*$^{-}$, derived from Miyagino | Itaya *et al*., 1999 |
| OK1 | *bio*$^{-}$, isolated from a commercial natto product | Kawamura, 1999 |
| Takahashi | *bio*$^{-}$, natto starter strain | Takahashi Yuzo Research Facility (Japan) |
| Naruse | *bio*$^{-}$, natto starter strain | Kubo *et al*., 2023 |
| *Bacillus licheniformis* JCM2505 | wild-type strain | RIKEN JCM (Japan) |
| *Bacillus megaterium* JCM2506 | wild-type strain | RIKEN JCM (Japan) |
| *Bacillus mojavensis* JCM12230 | wild-type strain | RIKEN JCM (Japan) |
| *Bacillus mycoides* JCM9801 | wild-type strain | RIKEN JCM (Japan) |
| *Bacillus pumilus* JCM 2508 | wild-type strain | RIKEN JCM (Japan) |
| *Bacillus velezensis* FZB42 | wild-type strain | *Bacillus* Genetic Stock Center (U.S.A.) |
| *Bacillus weihenstephanensis* KBAB4 | wild-type strain | Lapidus *et al*., 2008 |
| *Eschericha coli* | | Behnsen *et al*., 2013 |
| *Nissle 1917* | probiotic strain | |
| BL21 (DE3) SHuffle T7 Express lysY | MiniF, *lysY* / *fhuA2*, *lacZ*::*T7 gene1*, *lon*, *ompT*, *ahpC*, *gal*, *att*::pNEB3-r1-cDsbC (*lacI*$^{q}$), Δ*trxB*, *sulA11*, R(*mcr-73*::*miniTn10-Tet*$^{s}$)2, *dcm*, R(*zgb-210*::*Tn10-Tet*$^{s}$), *endA1*, Δ*gor*, Δ(*mcrC-mrr*)*114*::*IS10*; Cm$^{r}$, Spc$^{r}$ | NEB |
| *Yersinia pestis* Alexander | wild-type, virulent strain | Fukui *et al*., 1967 |
| **Plasmids** | | |
| pET22b(+) | IPTG-inducible protien expression vector; Amp$^{r}$ | Merck Millipore |
| pET-1×DT | pET22b(+) carring 3×FLAG–DogTag–His$_6$ tag | This study |
| pET-5×DT | pET22b(+) carring 3×FLAG–5×DogTag–His$_6$ tag | This study |
| pET-10×DT | pET22b(+) carring 3×FLAG–10×DogTag–His$_6$ tag | This study |
| pET-DC | pET22b(+) carring Dog Catcher–His$_6$ tag | This study |
| pET-mSc-DC | pET22b(+) carring mScarlet-I–Dog Catcher–His$_6$ tag | This study |
| pET-mCaf1-DC | pET22b(+) carring mCaf1–Dog Catcher–His$_6$ tag | This study |
| pET-mSc | pET22b(+) carring mScarlet-I–His$_6$ tag | This study |
| pET-mCaf1 | pET22b(+) carring mCaf1–His$_6$ tag | This study |

**Supplementary Table 2.**

Primers used in this study.

| Primer ID | Sequence (5′-3′)[a] |
|---|---|
| PKA-070 | ACAGTGTATAAGATCAATAC |
| PKA-009 | ggcccaaacgcatggCATGAGCGGATACATATTTG |
| PKA-010 | cggccatctccttggGTCTAGGGACCTCTTTAGCT |
| PKA-071 | ccatgcgtttgggccATCCCGCTTTAAGTCATCAC |
| PKA-072 | ccaaggagatggccgACTTCTCGTTGCGTTTGACG |
| PKA-073 | CATCGGCACCATTTTGTAGC |
| PKA-254 | ggcccaaacgcatggCAAAAATTATATGGAGATCT |
| PKA-255 | cggccatctccttggAAGCTTCACTAAATTAAAGT |
| PKA-313 | ccaaggagatggccgGTACTTCAATTATTACGTTA |
| PKA-314 | CCAAATTGTAATCGCCTTGT |
| PKA-739 | ccatgcgtttgggccTGTTTTGTTCCTCCCTGAAT |
| PKA-837 | TAGAGGTAAAGCAGTTAAAG |
| PKB-092 | TTAGCTCGAGCACCACCACCACCAC |
| PKB-093 | ATTTCATATGTATATCTCCTTCTT |
| PKB-134 | GGTGGTGGTTCAGGAGGAGGTAGC |
| PKB-134 | GGTGGTGGTTCAGGAGGAGGTAGC |
| PKB-134 | GGTGGTGGTTCAGGAGGAGGTAGC |
| PKB-137 | aaaactcgagCTTATAAAGTTCATCCATCC |
| PKB-138 | aaaaggatccGTATCTAAAGGTGAAGC |
| PKB-148 | cctcctcctgaaccaccaccCTTATAAAGTTCATCCATCC |
| PKB-310 | agccggcgatggccatggatACTCTTACATATAAGGAAGG |
| PKB-310 | agccggcgatggccatggatACTCTTACATATAAGGAAGG |
| PKB-311 | AAAGAAATCCTGGCTGCCCG |
| PKB-312 | cgggcagccaggatttctttGTTCGCTCAATTGGTTCCAAAG |
| PKB-313 | gccgttgccgctgcccgtTTGGTTAGATACGGTTAC |
| PKB-314 | aaacgggcagcggcaacggcGCAGATTTAACTGCAAGCAC |
| PKB-315 | tggtggtggtggtgctcgagGATGCGGGCTGGTTCAACAA |
| PKB-334 | agaaggagatatacatatgGACATTCCTGCGACATATGA |
| PKB-341 | tcctgaaccaccaccGATGCGGGCTGGTTCAACAA |
| PKB-345 | tggtggtggtggtgctcgagTTGCGGAACAATGCTTGTGA |
| PKB-345 | tggtggtggtggtgctcgagTTGCGGAACAATGCTTGTGA |
| PKB-345 | tggtggtggtggtgctcgagTTGCGGAACAATGCTTGTGA |
| PKB-345 | tggtggtggtggtgctcgagTTGCGGAACAATGCTTGTGA |
| PKB-352 | agaaggagatatacatatgGTATCTAAAGGTGAAGC |
| PKB-371 | GTGGTGGTGGTGGTGCTCGAGACATGAACCACCACCTTTTGGCG |
| PKB-405 | CACCACCACCACCACCACTGTTGAGATCCGGCTGCTAACAAAGCCCGA |
| PKB-406 | CTCGAGTGAACCACCACCTTTTGGCGGAATCGGTTC |
| PKB-478 | GATATTCCGGCTACGTATGAATTCACGGAT |
| PKB-479 | gatattgactacaaagatgatgacgataaaggatccGACATTCCTGCGACATAT |
| PKB-480 | gtgatctttatagtctccgtcgtgatctttataatcCATATGTATATCTCCTTC |
| PKB-481 | ggatccTTTATCGTCATCAT |
| PKB-492 | gcgcggatccGCTACCTCCTCCTGAACCACCACC |
| PKB-493 | ggccggatccGACATTCCTGCGACATATGA |
| PKB-563 | aagaaggagatatacatatgAAACTGGGCGAAATTGAG |

[a)] Engineered nucleotides are shown in lower letters.

### Supplementary Figures

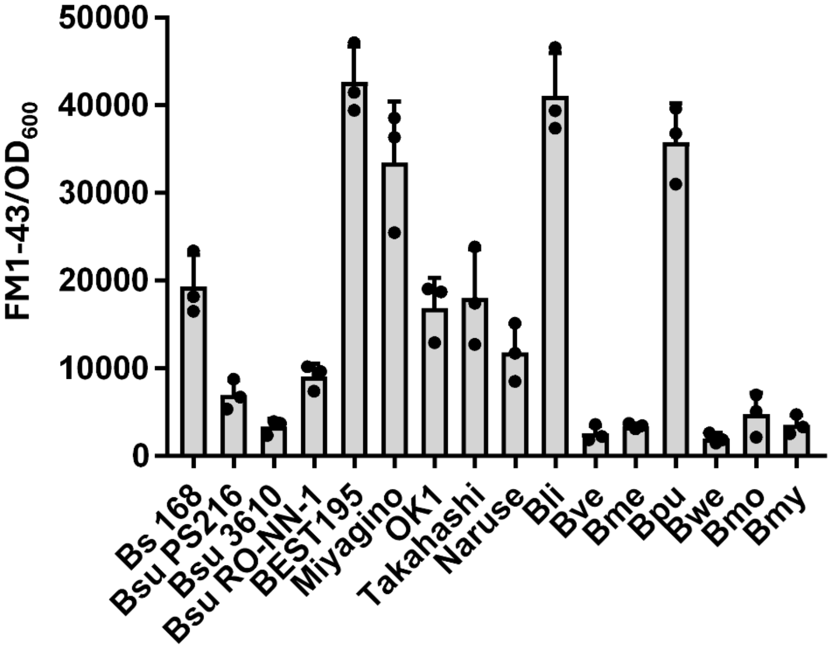

**Figure S1. MV productivity of *Bacillus* strains.** *Bacillus* strains were cultured in BHI medium for 24 h. MVs were isolated from the culture supernatants by ultracentrifugation. Relative amounts of the MVs were quantified using a lipid-specific fluorescent dye FM1-43. Error bars indicate ± standard deviations (SD) from three independent experiments. Bs 168, *Bacillus subtilis* 168; Bsu PS216, *B. subtilis* PS216; Bsu3610, *B. subtilis* NCIB3610; Bsu RO-NN-1, *B. subtilis* RO-NN-1; BEST195, *B. subtilis* Natto BEST195; Miyagino, *B. subtilis* Natto Miyagino; OK1, *B. subtilis* Natto OK1, Takahashi, *B. subtilis* Natto Takahashi; Naruse, *B. subtilis* Natto Naruse; Bli, *B. licheniformis* JCM2505; Bve, *B. velezensis* FZB42; Bme, *B. megaterium* JCM2506; Bpu, *B. pumilus* JCM2508; Bwe, *B. weihenstephanensis* KBAB4; Bmo, *B. mojavensis* JCM12230; Bmy, *B. mycoides* JCM9801.

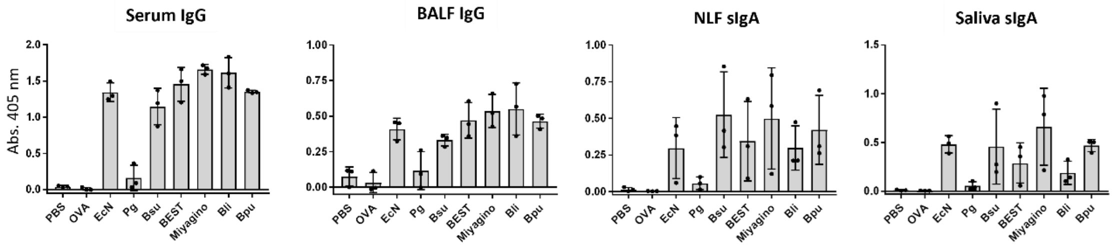

**Figure S2. Adjuvant activity of *Bacillus* MVs.** Mice were intranasally immunized three times at three-week intervals with admixtures of OVA and the indicated bacterial MVs. The production of OVA-specific antibodies was evaluated by ELISA. Error bars indicate ± standard deviations (SD) from three independent experiments. PBS, Phosphate-Buffered Saline; OVA, OVA alone; EcN, *E. coli* Nissle 1917 MVs; Bsu, BDM1 MVs (MV-producing host used in this study; derived from *B. subtilis* 168); Pg, *Porphyromonas gingivalis* MVs; BEST195, *B. subtilis* Natto BEST195 MVs; Miyagino, *B. subtilis* Natto Miyagino MVs; Bli, *B. licheniformis* JCM2505 MVs; Bpu, *B. pumilus* JCM2508 MVs.

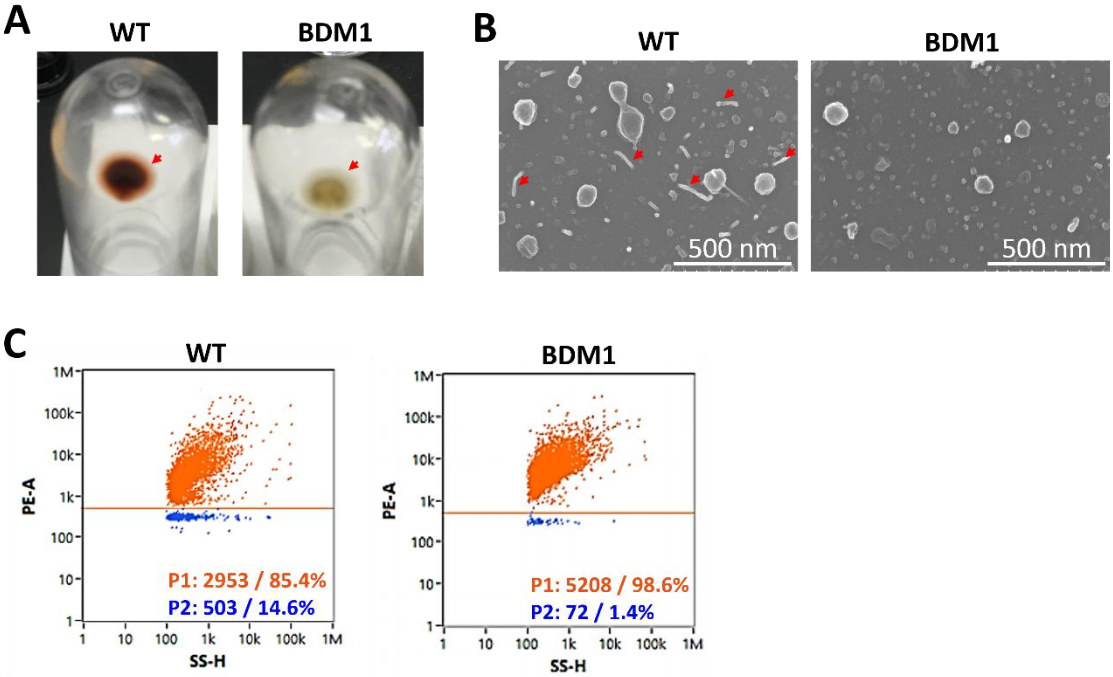

**Figure S3. Characterization of *B. subtilis* BDM1 MVs.** (**A**) **MV fractions of *B. subtilis* 168 WT and BDM1.** Culture supernatants of *B. subtilis* 168 WT and BDM1 (a Δ*hag* Δ*yvmC*–*cypX* strain derived from 168 WT) were ultracentrifuged to isolate MV fractions. The MV pellet (arrows) of 168 WT appears red-brown due to the presence of pulcherrimin. (**B**) SEM images. The MV fraction of *B. subtilis* 168 WT contains filamentous structures (arrows) due to the presence of flagella. (**C**) Nano Flow cytometry. MVs from *B. subtilis* 168 WT and BDM1 were labeled with the lipid-specific fluorescent dye FM1-43. The FM1-43-stained MVs were analyzed using NanoFCM Flare. Proportions (%) of FM1-43-positive particles (indicative of MVs; P1) and -negative particles (indicative of nonlipid particles including flagella; P2) are indicated in the panels.

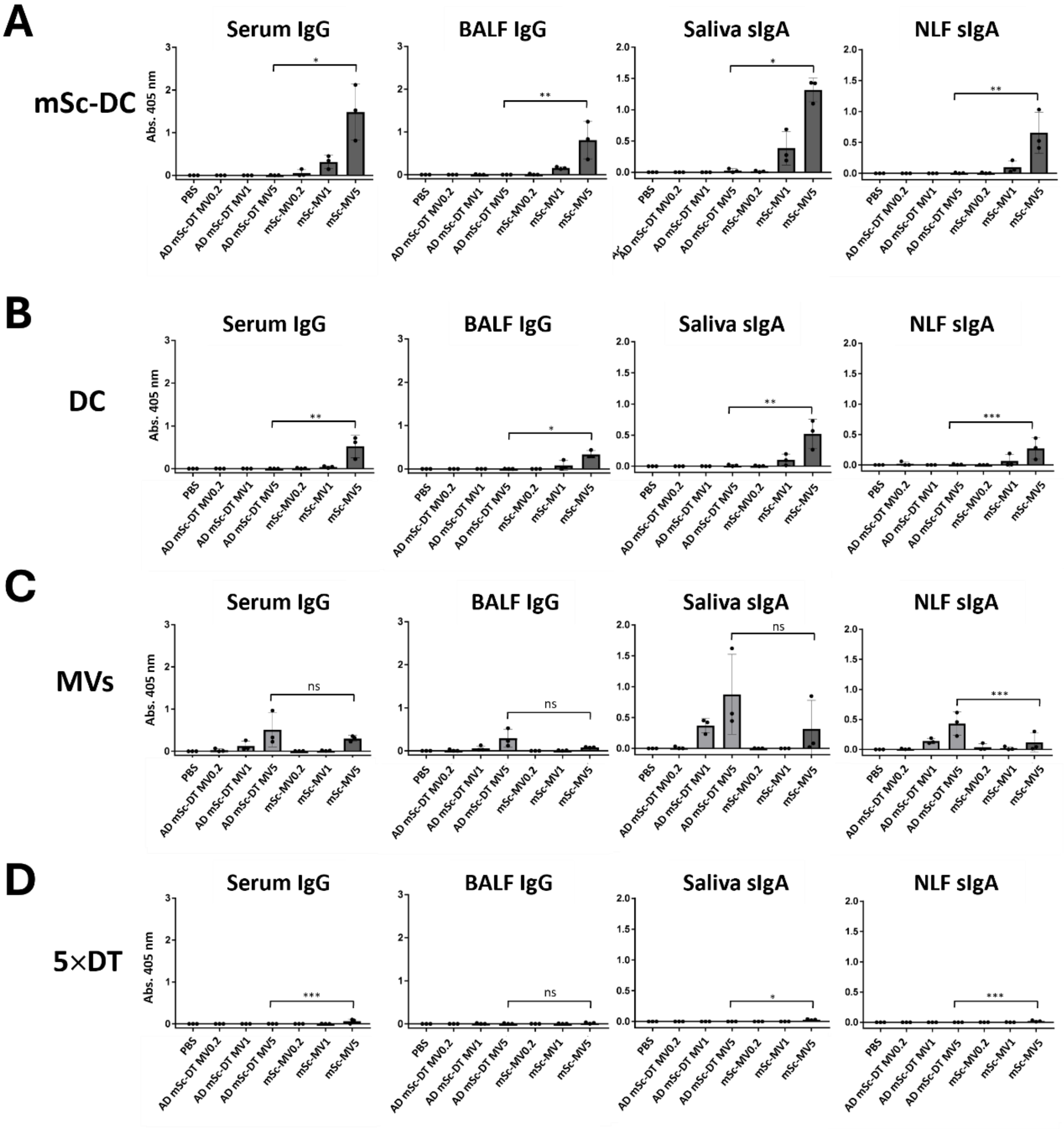

**Figure S4. Comparison of the immunogenicity between MV-anchored and unanchored multimerized mSc-DC.** Mice were intranasally administered three times at 2-week intervals with either the mSc-MV vaccine (mSc-MV0.2, 1, and 5) or an admixture of multimerized mSc-DC on 5×DT (AD mSc-DT MV0.2, 1, and 5). One week after the final immunization, ELISA was performed to assess production of IgG (in serum and BALF) and sIgA (in saliva and NLF) specific to mSc-DC (**A**), DC (**B**), MV backbone (**C**), and 5×DT (**D**). $n$ =3; ns, no significance; *$P$ <0.0001; **$P$ <0.001; ***$P$<0.025.

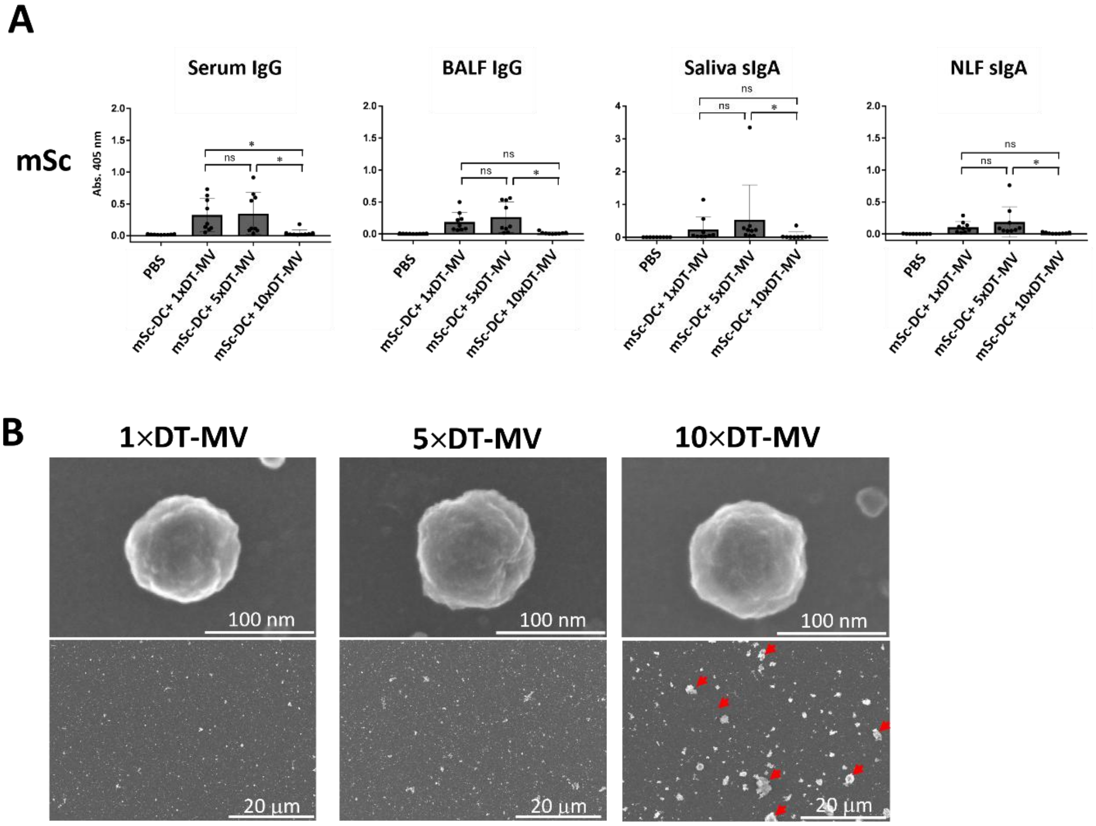

**Figure S5. Effects of the number of DT tandem repeats.** (**A**) Immunogenicity. Mice were intranasally administered mSc-MVs composed of mSc-DC and 1×, 5×, or 10×DT-MVs three times at 2-week intervals. Note that mSc-DC + 5×MV is identical to mSc-MV described in the main text. One week after the third shot, ELISA was performed to evaluate the production of IgG (in serum and BALF) and sIgA (in saliva and NLF) specific to mSc-DC (**A**). $n = 9$; ns, no significance; $*P < 0.05$. (**B**) SEM images. Tz-MVs were conjugated with either 1×, 5×, or 10×DT scaffolds and observed using SEM. Arrows indicate aggregated 10×DT-MVs.

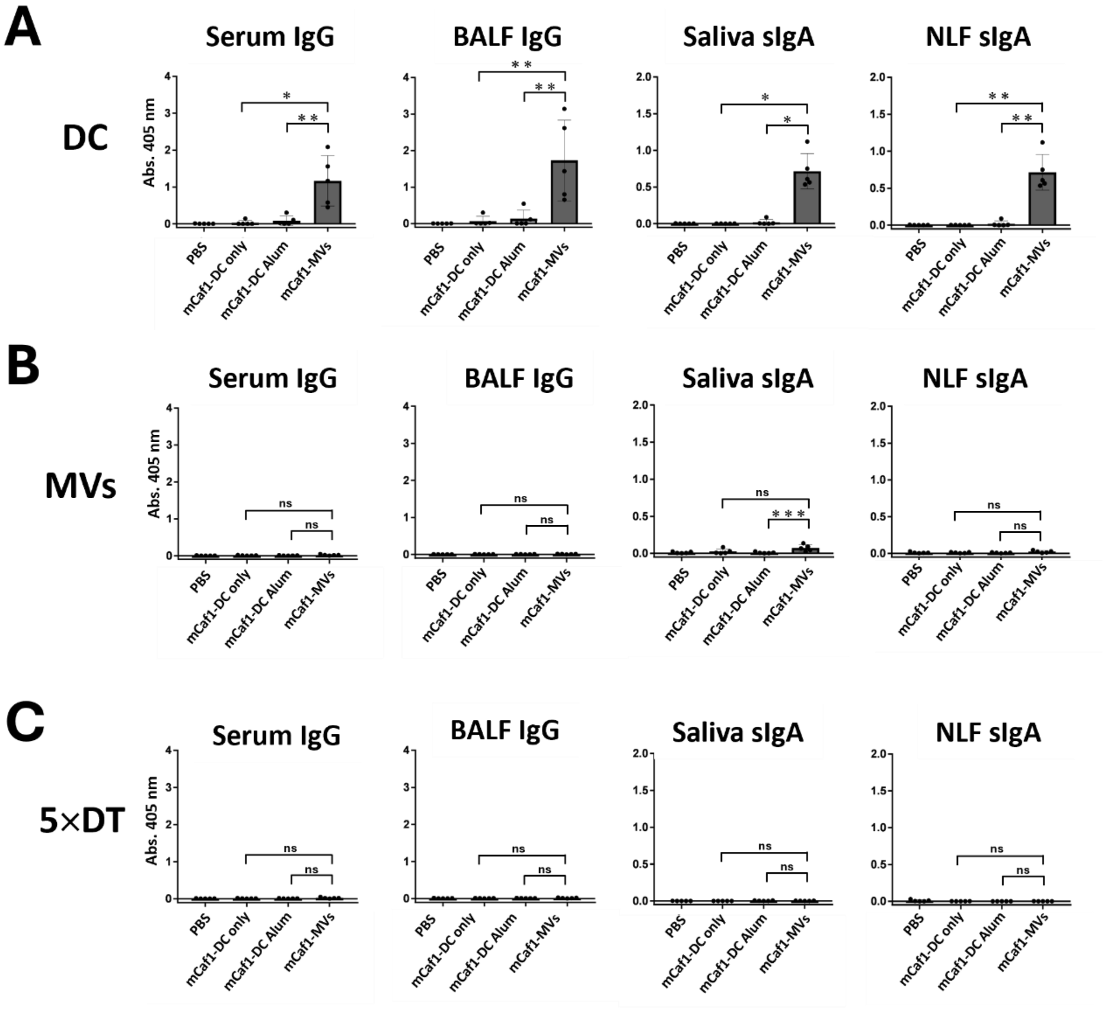

**Figure S6. Immunogenicity of *Y. pestis* vaccines.** ELISA was performed to evaluate the production of IgG (in serum and BALF) and sIgA (in saliva and NLF) specific to DC (**A**), the MV backbone (**B**), and 5×DT (**C**). Data regarding mSc-specific antibodies are presented in the main text and Figure 5. $n$ = 5; ns, no significance; *$P$ <0.0005; **$P$ <0.005; ***$P$<0.05.

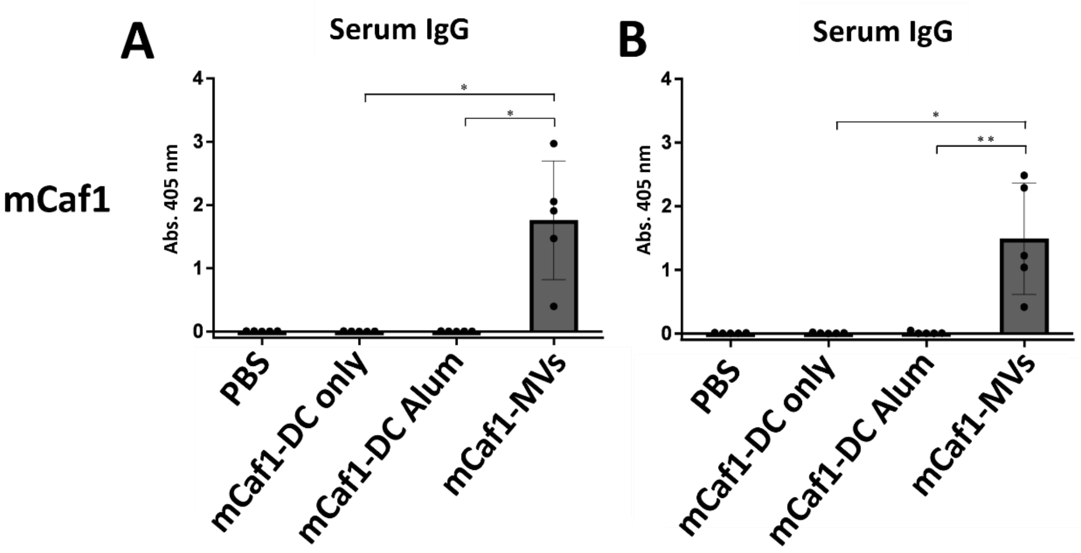

**Figure S7. Confirmation of IgG production prior to *Y. pestis* challenge.** Levels of the mCaf1-specific IgG antibodies in sera from mice immunized three times were confirmed prior to the *Y. pestis* infection experiment (shown in Figure 6). Mice in panels **A** and **B** were subjected to 10× and 100× $LD_{50}$ infections, respectively. $n = 5$; $*P = 0.0001$; $**P = 0.0005$.
